# *Andrographis saxatilis* (Acanthaceae) a new woody species from the Eastern Ghats of Andhra Pradesh, India

**DOI:** 10.1101/2023.03.19.533309

**Authors:** Raja Kullayiswamy Kusom, Sarojini Devi Naidu

## Abstract

*Andrographis saxatilis* (Acanthaceae) a woody undershrub, is described and illustrated from the rocky plateaus of the Owk dam area, Andhra Pradesh. It resembles *A. beddomei* but differs in many morphological characters.

## INTRODUCTION

*Andrographis* Wall ex Nees is a tropical Asian genus (Mabberley, 2008) of 58 taxa and 44 species (POWO, 2023), with 29 taxa occurring in India (Karthikeyan et al., 2009, Gnanasekaran et al., 2016), especially Peninsular India of which 23 are endemic (Gnanasekaran & Murthy, 2012; Gnanasekaran et al., 2015). *Andrographis* is well known by *A. paniculata* which is most popular in Ayurvedha medicine for its anticancer, antimicrobial, antifungal, antihelmenthic, etc. properties. (Okhuarobo et al 2014). *Andrographi serpilifolia* (Venkata et al 2021), *A. liniata* (Bhat and Hoskatte 2023) also resembles *A. paniculata* in medicinal properties. *Andrographis* species are mostly herbaceous and annuals but some are perennial with a stout stem or/and root (*Andrographis beddomei* and the present described new species). Insufficient study has been done on this genus. Understudying more about this group of species from an evolutionary point of view is very important.

While doing a field survey under the Dharmavana Nature Ark (DNA) Peninsular India project, the authors collected an interesting woody *Andrographis* species. It is a perennial undershrub that grows between red lime stone with a stout stem in scrub forest near the Owk dam area. On critical examination it clearly showed differences compared to the known species of the genus. It is described and illustrated herein with colour photographs and comparative characteristics with allied species for easy identification.

## MATERIAL AND METHODS

Collected specimens were treated with 70% ethanol and made herbarium specimens following standard methods (Santapau 1958, Jain and Rao 1977). Flowers and anthers were fixed in 70% ethanol for future reference. Morphological and microscopic characters were studied under Olympus SZ61 with Magnus (Magcam DC 5) camera attached.

### TAXONOMIC TREATMENT

Family Acanthaceae Juss.

Genus *Andrographis* Wall. ex Nees.

**Andrographis saxatilis** Raja Kullayisw. & Sarojin. sp. nov. (Fig. 1, 2; Table 1).

**FIG. 1.** — *Andrographis saxatilis* Raja Kullayisw. & Sarojin., sp. nova (drawing by N. Sarojini Devi): **A**, flowering branch; **B**, lower surface of leaf; **C**, upper surface of leaf with petiole; **D**, flower; **E**, corolla split open; **F**, gynoecium; **G**, capsule; **H**, calyx; **I**, dehised fruit; **J**, seed. Scale bars: A, 1.5 cm; B, 3 mm; C, H, J, 1 mm; D, 3 mm; E, 1.5 mm; F, 2.6 mm; G, I, 2.5 mm.

**FIG. 2.**
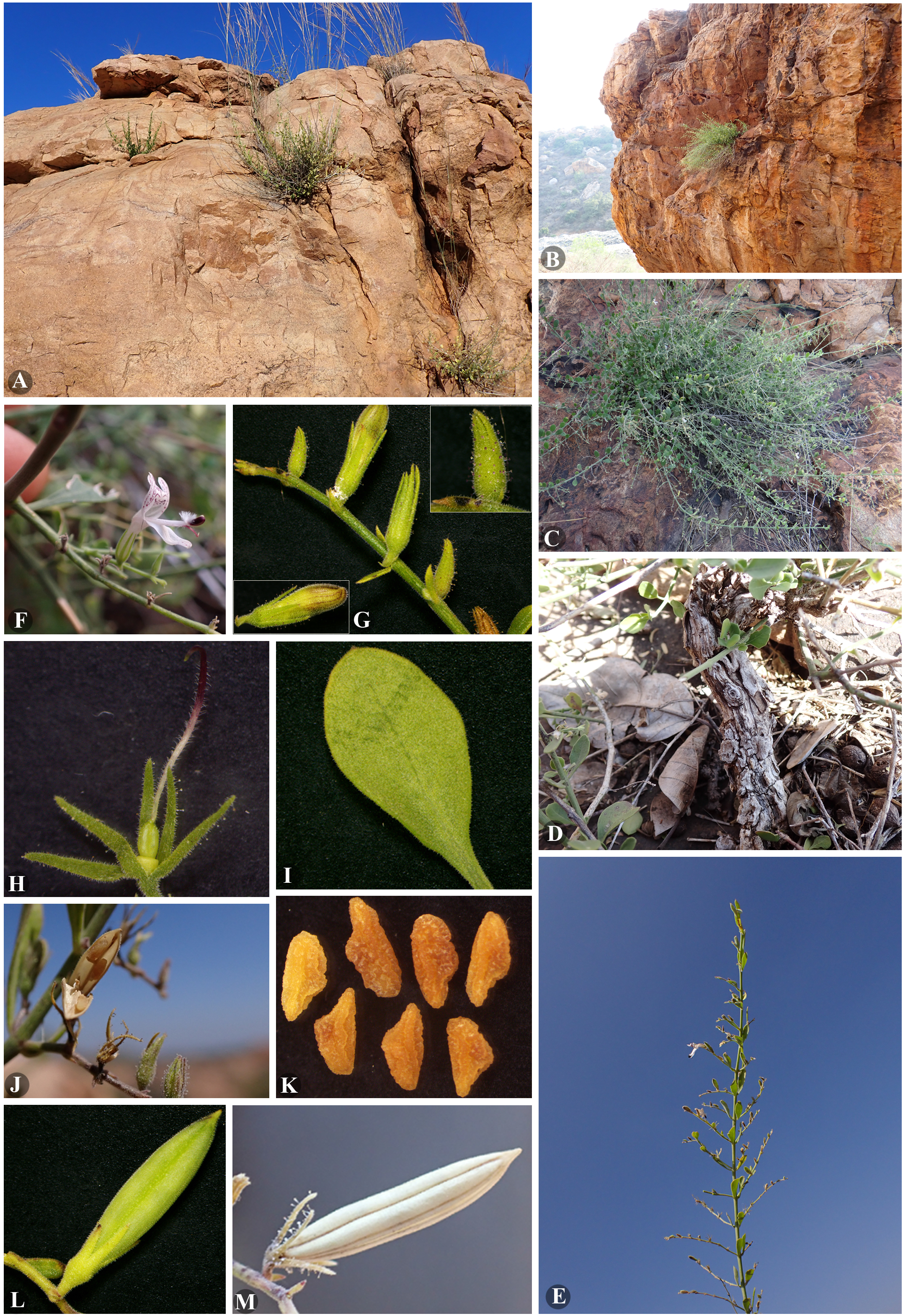
*Andrographis saxatilis* sp. nova. **A**&**B**, habitat; **C**, habit; **D**, stout stem; **E**, twig; **F**, flower; **G**, flower buds and calyx; **H**, calyx along with ovary and style; **I**, leaf; **J**, open fruit along with seeds; **K**, seeds; **L**, young fruit; **M**, mature fruit.

**Table 1:** Comparative table of closely related species

| Characters | *Andrographis saxatilis*<br>**sp. nov.** | *A. beddomei* | *A. paniculata* |
| --- | --- | --- | --- |
| Habit | Rocky crevices | Hard gravelly soil | Gravelly and loose soil |
| Root | Woody, stout | Tuberous woody. stout | Normal tap root |
| Stem | Woody and stout | Herbaceous | Herbaceous |
| Branches | Erect, rarely decumbent,<br>basally 4-angular at base<br>only | Decumbent to prostrate, 4-angular | Erect, 4-angular |
| Leaves | Obovate, elliptic-oblong,<br>rounded apex | Obovate to narrowly<br>oblanceolate, elliptic,<br>rounded apex | Elliptic-oblong,<br>acuminate apex |
| Venation | Obscure | Prominent | Prominent |
| Inflorescence | Axillary, larger than the<br>leaves, unbranched | Axillary and terminal,<br>shorter than the leaves,<br>unbranched | Axillary and terminal,<br>larger than the leaves,<br>branched |
| Flowers | 3 – 8 flowers, glandular<br>hairy on sepals, outside<br>of corolla tube | 2 – 3 flowers, glandular<br>hairy on sepals, out and<br>inner side of corolla tube | 8 – 15 flowers, glandular<br>hairy on sepals, outside<br>of corolla tube |
| Stamens | Larger than the lower lip | Shorter or as long as | Larger than the lower lip |
|  | of corolla | lower lip of corolla | of corolla |
| Lower lip of corolla | Unlobed | Slightly lobed | Lobed |
| Pedicels | 1 – 2 mm long | 4 – 15 mm long | 8 – 17 mm long |
| Fruit | Non glandular hairy | Glandular hairy | Glandular hairy |

*Andrographis saxatilis* sp. nov. resembles *A. beddomei* by its stout root, obovate leaves, glandular calyx, pubescent corolla, seed colour and shape, but it is distinguishable from the later by its growth pattern, smaller leaves, racemes larger than leaves, cylindrical stem, glaucous green stem and leaves, smaller pedicel, stamens that are larger than the lower lip of corolla, nonglandular hairs on fruit and some more characters that are tabulated in table 1.

### Type material

**India**. Andhra Pradesh, Nandyal district, Owk mandal, Owk dam area, rocky plateaus 15.227718N, 78.082321E, alt. *c*. 282 m, 3.I.2023, *Raja KSwamy* DNA001 (holo- BSID! · iso- MH!, SKU!, HDNA!).

### Etymology

The species epithet “*saxatilis*” describes the habitat rocky crevices of the plants.

### Distribution

*Andrographis saxatilis* sp. nov. is known only from Owk rocky plateaus, near Owk reservoir, about 8 sq. km area., Nandhyala district, Andhra Pradesh, South India.

### Habitat

Rocky crevices of scrub forest at an altitude 270 – 290 m. P

### henology

Flowering fruiting from December to March.

### Ecology

Dry red gravelly soil, between red sandy rocks at an elevation of 270 – 282 m a.s.l. Associated with *Aristida setacea* Trin., *Heteropogon contortus* (L.) P.Beauv. ex Roem. & Schult., *Pseudopogonatherum trispicatum* (Schult.) Ohwi, *Phyllanthus palakondensis* Raja Kullayisw. & Sarojin.

### Conservation status

Population size of this species is 458 individuals per 8 sq. km area. It is not found in entire rocky plateaus of the same location which spreads around a 279 sq. km, but found only in 8 sq. km area. This species has been categorised under data deficiency (DD) as there was insufficient information about its occurrence and distribution in India. It is evaluated here as ‘Vulnerable’[VU Bab(iii)+2ab(iii)] using the IUCN Red List Categories and Criteria version 15.1 (IUCN 2022). Main threats observed were grazing and forest fire. Fire damages like branches along with flowers and fruits mostly when these plants associate with *Heteropogon* and other grass. We also observed that seed setting is very poor, it might be due to a lack of pollinator or sexual incompatibility. About 500 fruits were collected in three periodical field visits, among them 8 fruits were with fully developed seeds (8 seeds per capsule), about 10 fruits with 1 – 2 developed seeds and the rest were abortive. The potted plant in DNA nursery has fruits but without seed development, probably due to lack of cross pollination.

### Additional Specimens Examined

**India**. Andhra Pradesh, Nandyal district, Owk mandal, Owk dam area, rocky plateaus 15.228514N, 78.081039E, alt. *c*. 276 m, 8.I.2023, *Raja KSwamy* DNA115 (HDNA!, BSID!, MH!, SKU!). — **India**. Andhra Pradesh, Nandyal district, Owk mandal, Owk dam area, rocky plateaus 15.229095N, 78.079533E, alt. *c*. 278 m, 6.III.2023, *Raja KSwamy* DNA142 (HDNA!).

## DESCRIPTION

Perennial woody undershrub, up to 50 cm, stem stout, bark grey, fissured vertically. Branches many from woody root stock, basally slightly angular, hispid and terete towards apices, pubescent, glaucous green, corky, 2 – 3 mm in diameter. Leaves opposite decussate, obovate, cuneate, elliptic-oblong, 8 – 25 × 5 – 11 mm, pubescent, glaucous green, slightly coriaceous, leaf base tapering into petiole, margin entire, apex rounded, mid vein inconspicuous above, prominent below; lateral veins 3 pairs, alternate, obscure above, prominent below; petiole 3 – 5 mm long, pubescent. Inflorescence axillary, raceme; rachis 3 – 8 cm long, pubescent, terete, unbranched, 3 – 8-flowered; flowers distinctly arranged (interstices 4 – 8 mm long, reduces at apices); peduncles 1 – 1.5 cm long, pubescent; pedicel 1 – 1.5 mm long. Bracts foliaceous, spathulate, mucronate, pubescent, 4 × 2 mm base of rachis and reduces towards apices; bracteoles 2, subulate, lanceolate, 1 – 1.5 mm long, pubescent. Flowers 1.5 – 1.7 cm long, 0.4 – 0.5 cm across. Calyx green, 6 – 7 mm long; sepals 5, linear-lanceolate, acuminate at apex, glabrous within, glandular pubescent without. Corolla white, bilipped; tube 5 – 6.5 mm long, densely villous inside, sparsely glandular pubescent outside; mouth of the tube swollen, 4 – 5 × 5 – 6 mm, glandular pubescent within and without; lower lip of corolla boat shaped at young, 9 – 10 × 2 – 3 mm, ligulate and curved back at later stages; upper lip connate part 3.8 – 4.2 × 5 – 5.5 mm; lobes 3, ovate, 3 – 3.5 × 1.6 – 2 mm with dark purple-striped, margin entire, apex obtuse, glabrous within, glandular pubescent without, middle lobe hirsute at centre, bent back at maturity. Stamens 2, adnate at base of ventricose portion of corolla tube; filaments 10 – 13 mm long, white, sparsely hispid, dilated at base, retrorsely villous at attachment; anthers linear to oblong, dark purplish, adnate, 2.3 – 3 × 2.1 – 3 mm together, woolly tomentose at base; each anther oblong, 2.3 × 1.5 mm. Ovary ovoid-oblongoid, 1.5 – 2 × 1 mm, bristled hispid, receptacle swollen, white, 1.5 mm across. Style 13 – 14 mm, antrorsely bristled hirsute; stigma linear, curved, deep purple, green at tip. Capsules linear-oblong to very narrowly ellipsoid, acute at ends, 15 – 17 × 3 – 4 mm, tomentellous, 4 – 8-seeded. Seeds yellowish, sub-qudrate, 2.5 – 3 × 1.5 mm, oblique at base, obtuse at apex, hard, scrobiculate, and verrucose.

## Key to the *andrographis* ALLIED SPECIES

1. Stems and roots stout and woody, leaves obovate, base narrowed …………….2 −Stems and roots not as above, leaves ovate-oblong or elliptic, base acute ……………3
2. Branches decumbent, angular, glabrous, inflorescences shorter than the leaves, leaves 2.5 – 9 cm long ……………………………………………….*A. beddomei* C.B.Clarke −Branches erect (rarely decumbent), terete, hispid hairy, inflorescence larger than the leaves, leaves 0.8 – 2.5 cm long …………………………….*A. saxatilis* Raja Kullayisw. & Sarojin. sp. nov.
3. Leaves narrow, ovate-oblong to lanceolate, glabrous ………..*A. paniculata* Wall. ex Nees −Leaves ovate to orbicular, glandular hairy ……………………………..*A. rothii* C.B.Clarke

## Acknowledgements

The authors are grateful to the DNA for facilities and making periodical field visits for gathering more data on the species distribution and population size, and also thank to the DNA jungle team for supporting. This species is proposed for ex-situ conservation within the Dharmavana nature ark.

## Legends

Map: Distribution map of *Andrographis saxatilis* and allied species

